# Insights into β-manno-oligosaccharide uptake and metabolism in *Bifidobacterium adolescentis* DSMZ 20083 from whole-genome microarray analysis

**DOI:** 10.1101/2022.03.08.483560

**Authors:** Priyanka Rose Mary, P Monica, Mukesh Kapoor

## Abstract

**Aims:** To determine the ability of adult-associated *B. adolescentis* DSMZ 20083 to utilize dietary β-manno-oligosaccharides and understand the underlying molecular mechanism.

**Methods and Results:** *In-vitro* fermentation and TLC were used to determine the ability of *B. adolescentis* DSMZ 20083 to utilize β-manno-oligosaccharides from guar gum, locust bean gum, konjac and copra meal. Further, Whole-genome transcriptome analysis, q-RT-PCR and molecular docking were employed to reconstruct copra meal β-manno-oligosaccharides (CM-β-MOS) utilization pathway in *B. adolescentis* DSMZ 20083.

*B. adolescentis* DSMZ 20083 grew appreciably (A_600nm_ up to 0.8) on all tested β-manno-oligosaccharides but maximally on CM-β-MOS. Transcriptome analysis revealed differential up-regulation of three distinct gene clusters encoding, ABC transporter cassette (ESBP’s and permeases), MFS transporter, GH1 β-glucosidase and, GH32 β-fructofuranosidases. ABC and MFS possibly transported majorly DP≥2 and DP≥3 CM-β-MOS, respectively. Sugar absorption and utilization pathways; ABC transport system, pyruvate metabolism, glycolysis/gluconeogenesis, pentose, and glucouronate inter-conversions were also up-regulated.

**Conclusions:** *B. adolescentis* DSMZ 20083 possibly utilizes ABC & MFS transporters to internalise and GH1 β-glucosidase, and GH32 β-fructofuranosidase to deconstruct CM-β-MOS.

**Significance and Impact of the Study:** First study reporting possible molecular determinants used by *B. adolescentis* DSMZ 20083 to utilize β-manno-oligosaccharides and thus, can prove resourceful in precision microbiome modulation.

## 1. Introduction

Dense and diverse human gut microbiota (HGM) inhabiting the gastrointestinal tract is a pivotal determinant in maintaining host health (Sonnenberg and Backhed, 2016). The orchestrated breakdown of recalcitrant dietary glycans such as mannan, xylan, xyloglucan, β-glucan, pectin, and resistant starch by a repertoire of carbohydrate-active enzymes (CAZymes) and associated non-catalytic proteins, contributed by HGM plays a key role in modulating its architecture and metabolic output (Ndeh and Gilbert 2018; Tamura and Brumer 2021).

β-mannans are homo- or hetero-polymers of β-1,4 linked mannose residues which are interrupted with glucosyl moieties in glucomannan, or substituted by galactosyl residues in galactomannan and are at times acetylated, as in glucomannan or galactoglucomannan. They occur as structural components in the plant cell wall and energy reservoir in the endospermic component of seeds and nuts such as locust bean, guar seeds, coconut palm, konjac root and coffee bean (Srivastava and Kapoor 2017; Srivastava *et al*. 2017; Mary *et al*. 2019). Besides being a source of naturally occurring dietary fibre, they are widely employed as hydrocolloids in various food products like sauces, ice-creams, yogurts, and fruit juices thereby making them an integral component of the present-day human diet (Srivastava and Kapoor 2017). β-manno-oligosaccharides generated from various β-mannans have been shown to selectively favour the growth of beneficial gut bacteria (Srivastava *et al*. 2017; Mary *et al*. 2019; Panwar and Kapoor 2019). A growing body of evidence suggests that consumption of β-mannans or β-manno-oligosaccharides leads to improved intestinal barrier function, glucose homeostasis, immune stimulation, lowered serum lipids, prevention of colon carcinogenesis, osteoporosis, and obesity which is consistent with their utilization by the human gut bacteria (La Rosa *et al*. 2019; Jana *et al*. 2020; Gill *et al*. 2021).

The genus *Bifidobacterium* belonging to the phylum Actinobacteria are G+C rich, Gram-positive bacteria (Milani *et al*. 2017). Members of this genus are known to improve bowel function, safeguard from inflammatory disorders, colorectal cancer (in animal models), and enteropathogenic infections (Fukuda *et al*. 2011; O’Callaghan and Van Sinderen 2016). Certain members of this taxon such as *Bifidobacterium longum* subsp *infantis, Bifidobacterium breve*, and *Bifidobacterium bifidum* are more prominent in breast-fed infants (Turroni *et al*. 2018). However, post-weaning when there is shift from a milk-based diet to a plant-based diet, the taxon is represented by adult species such as *Bifidobacterium adolescentis*, *Bifidobacterium cataneulatum*, and *Bifidobacterium longum* subsp. *longum*. This change in species composition is due to their saccharolytic preferences biased towards plant-derived carbohydrates (Palmer *et al*. 2007; Koenig *et al*. 2011).

Bifidobacteria are considered secondary utilizers but are capable of competitive growth with generalist degraders like *Bacteroides ovatus* and *Roseburia intestanilis* as their utilization patterns are skewed towards oligosaccharides by virtue of their arsenal of ABC transporters (Ejby *et al*. 2016; Bågenholm *et al*. 2017; Turroni *et al*. 2018; La Rosa *et al*. 2019; Ejby *et al*. 2019; Fushinobu and Hachem 2021). *B. adolescentis* ATCC15703 has been recently shown to ferment β-manno-oligosaccharides from cassia gum, Norwegian spruce wood, and low viscosity LBG (La Rosa *et al*. 2019; Li *et al*. 2020; Bhattacharya *et al*. 2021) but the molecular determinants responsible for utilization remains obscure as specific transporters and glycoside hydrolases are yet to be elucidated.

In the present study, we report utilization of β-mannooligosaccharides derived from copra meal, guar gum, locust bean gum, and konjac β-mannans by *B. adolescentis* DSMZ20083/ATCC15703. Further, using a whole-genome transcriptome analysis complemented by real-time PCR studies, the gene clusters possibly involved in the uptake and catabolism of β-manno-oligosaccharide in *B. adolescentis* DSMZ20083 were identified for the first time. Insights into the specificity and binding affinity of the up-regulated extracellular solute binding protein (bglE) towards β-manno-oligosaccharide were gained through computational molecular docking.

## 2. Material and methods

### 2.1 Carbon sources

Polymeric dietary β-mannans; locust bean gum, konjac glucomannan, and guar gum were procured from Sigma Aldrich, USA; Megazyme, Ireland and Hi-media laboratories, India, respectively. Copra meal prepared from dried and desiccated coconut flour (Prabha *et al*. 2022 under review) was procured from the laboratory. Commercial β-manno-oligosaccharides; mannobiose (CM2), mannotriose (CM3), and mannotetrose (CM4) were purchased from Megazyme, Ireland. Crude short-chain dietary β-manno-oligosaccharides derived from locust bean gum (GMOS, Srivastava *et al*. 2017), guar gum (GG-β-MOS) and konjac glucomannan (KGM-β-MOS) (Mary *et al*. 2019), and copra meal [CM-β-MOS, Prabha *et al*. 2022 (under review)] for use in *in-vitro* fermentation studies were prepared using GH26 endo-mannanase; ManB-1601. To avoid any batch-wise variations, β-manno-oligosaccharides produced in a single batch were used to carry out all the experiments. The monosaccharides used in the present study were glucose (Glc), galactose (Gal), and mannose (Man) and procured from Sigma-Aldrich (MO, USA).

### 2.2 Bacterial strain and fermentation media

*Bifidobacterium adolescentis* DSMZ 20083 [Reuter E194a (Variant a) isolated from human intestine] was procured from DSMZ, Germany. The strain was routinely cultured (static) at 37 °C under anaerobic conditions (Anaerobic gas jar, Hi-Media, India) in deMan, Rogosa, Sharpe broth (MRS, Hi-Media, India). For the *in vitro* growth assays, culturing was done in <u>synthetic defined MRS medium</u> (SDM, w/v or v/v) comprising of protease peptone 1%, beef extract 1%, yeast extract 0.5%, ammonium citrate 0.2%, sodium acetate 0.5%, MgSO_4_ 0.01%, MnSO_4_ 0.005%, K_2_HPO_4_ 0.2% and tween-80 0.1 ml, supplemented with 0.05 % cystine-HCl or <u>minimal media</u> (MM, w/v or v/v) prepared in the following manner: Yeast extract (2.5 g/l), tween 80 (1 % v/v) Nacl (0.2 g/l), KCl (0.25 g/l), NH_4_Cl (0.40 g/l) were dissolved in double distilled water and autoclaved (121 °C, 20 min). Prior to use, a mixture of 0.15 g/l MgCl_2_.6H_2_O, 6 g/l KH_2_PO_4_, 4 g/l Na_2_HPO_4_ and 0.5 g/l cystine-HCl was filter-sterilized (0.2 μm) and added to the sterilized media. All the media components used in the present study were of analytical grade and procured from either Hi-media laboratories, India, or Merck, India.

### 2.3 *In vitro* fermentation studies

The ability of *B. adolescentis* DSMZ 20083 to grow on polymeric dietary β-mannans (locust bean gum, guar gum, konjac and copra meal), commercial β-manno-oligosaccharides (CM2, CM3 & CM4), and dietary β-manno-oligosaccharides (GMOS, GG-β-MOS, KGM-β-MOS, and CM-β-MOS) was assessed by *in vitro* fermentation in SDM. The media was supplemented with respective filtered-sterilized (0.22 μm) carbon source (0.5 % w/v; polysaccharides, commercial β-manno-oligosaccharides or dietary β-manno-oligosaccharides) and inoculated with 1 % (v/v) of primary inoculum (20 h-old culture of *B. adolescentis* DSMZ 20083 grown in MRS media). Thereafter, the cultures were incubated in an anaerobic gas jar at 37 °C for 48 h. The growth of *B. adolescentis* DSMZ 20083 in the fermentation broth was estimated by measuring the O.D_600nm_, log CFU ml^−1^ and pH after 48 h. For the measurement of log CFU, cultures were serially diluted and plated onto MRS agar supplemented with cystine-HCl (0.05 % w/v) and incubated at 37 °C in an anaerobic gas box for 48 h. The growth curves of *B. adolescentis* DSMZ 20083 on CM-β-MOS, monosaccharide controls (Glc and Gal), and carbon-free media were performed under the culture conditions described above for up to 72 h. Samples were aliquoted at regular intervals to estimate the cell growth (O.D_600nm_). Generation time was calculated by measuring the rate of increase in the O.D_600nm_ values between 0.2 and 0.6.

The depletion of dietary β-manno-oligosaccharides [GMOS, GG-β-MOS and CM-β-MOS, predominant species ≥ 75 % DP2 and DP3 confirmed by ultra-performance liquid chromatography electron spray light scattering detector (UPLC-ELSD, Waters, USA) (Mary *et al*. 2019)] and monosaccharides (Glc, Gal and Man) from the growth media was studied using TLC as mentioned previously by Rogowski *et al*. (2015) with certain modifications. Fermentation was carried out in MM supplemented with the carbon sources [Glc, Gal, Man, GMOS, and CM-β-MOS were added at a concentration of 0.5 % (w/v) while, GG-β-MOS was supplemented only at 0.1 % w/v (A_600nm_ of *B. adolescentis* DSMZ 20083 was found to decrease with increase in the concentration of oligosaccharides to 0.25 and 0.5 % w/v; data not shown)] and the broth was withdrawn at appropriate intervals and analysed for un-utilized oligosaccharides left in the broth. An aliquot of 5 μl was spotted on an activated (100 °C, 1 h) TLC sheet (Merck, USA). CM2, CM3, and CM4 were included as detection standards. Plates were clamped on top with Whatman filter paper to absorb the excess solvent and were resolved over-night (12-15 h) using butanol: water: acetic acid (2:1:1) as the mobile phase. Thereafter, plates were air-dried and developed with a mixture of orcinol (1 mg/ml), ethanol (75 % v/v), and sulphuric acid (3.2 % v/v) prepared in 100 ml of Milli Q water followed by heating the plate at 100 °C for 10 min.

### 2.4 Transcriptional analysis

*B. adolescentis* DSMZ 20083 was grown till mid-log phase (~20 h) in MM supplemented with Glc (0.5 % w/v) and was inoculated into MM supplemented with CM-β-MOS (0.5 % w/v) or control sugar (Glc) separately and passaged three times in the respective carbohydrate supplemented media by incubating anaerobically at 37 °C until O.D_600nm_ reached 0.8 (late log phase; ~36 h incubation). For the final passage (3^rd^) on CM-β-MOS, enrichment of short-chain fractions (DP2-5) was carried out by subjecting crude CM-β-MOS to purification on Bio-gel P2 (Prabaha *et al*. 2022 under review). The purified peaks having DP2-5 were pooled, freeze-dried, and supplemented into the growth medium. The bacterial cells from the 3^rd^ passage were collected by centrifugation (5000x g, 10 min) and the pellet obtained was washed twice (8.500x g, 10 min) using phosphate buffer saline (prepared in nuclease-free water). The pellet was then re-suspended in 5 ml of RNA later (Sigma Aldrich, MO, USA) and incubated overnight at 4°C. Thereafter, the re-suspended pellet was stored at −80°C until RNA was isolated. The total RNA was extracted using Trizol reagent +Rnaeasy Mini Kit (Qiagen, Germany) by following the supplier’s protocol. The concentration and quality of purified RNA were determined by using nano-drop (ND 2000, Thermo Scientific, USA) and bio-analyzer (Agilent 2100, Bio-analyzer, USA), respectively. 16S and 23S rRNA ratio and RNA integrity number were calculated from 2100 Expert software and RIN Beta Version Software (Agilent), respectively. The purified RNA was stored at −80°C until further usage.

A custom whole-genome microarray (Genotypic Technology Private Limited, Bengaluru, India; 8×15K: AMADID: 86602) of 1661 genes for *B. adolescentis* DSMZ 20083 was designed with an average of 3 replicate spots for each gene. cRNA generation, fragmentation, and hybridization were performed as described previously (Panwar and Kapoor 2019). The hybridized slides were washed using gene expression wash buffer (Agilent, USA) and scanned. Microarray images were collected by using sure scan microarray scanner system at 5 μm resolution (Agilent Technologies, Scanner 2600D). Fluorescent intensities were quantified using feature extraction software. Gene Spring GX software (Agilent Technologies) was used to normalize the raw data using the percentile shift normalization method. In each sample, expression values were obtained by subtracting the log_2_ transformed intensity values for each probe by the calculated 75^th^ percentile value of the respective array. Genes showing fold change > 1 and fold change < −1 were considered up-regulated and down-regulated, respectively in the test, with respect to controls. A volcano plot algorithm was used to calculate statistical student t-test and *p* value among the replicates. In order to identify significant gene expression patterns, hierarchical clustering of differentially regulated genes was done on the basis of Pearson coefficient correlation algorithm. Details of the microarray platform are available at http://www.ncbi.nlm.nih.gov/geo under the accession number GSE180349.

### 2.5 Real-time quantitative PCR (q-RT-PCR)

Ten differentially expressed genes [BAD_RS06080, BAD_RS06835, BAD_RS06830, BAD_RS06825, BAD_RS06820, BAD_RS06085, BAD_RS06090, BAD_RS05480, BAD_RS02080, and BAD_RS05750] from the microarray analysis were selected for real-time quantitative PCR in order to validate microarray data. Primer sequences (Supplementary Table 1) for genes were designed and analyzed using the Primer3 software. *B. adolescentis* DSMZ 20083 was first passaged in MM supplemented (0.5 % w/v) with either Glc or CM-β-MOS as the sole carbohydrate source by incubating at 37 °C under anaerobic conditions for 36 h (O.D_600nm_ ~0.8). Thereafter, a second passage (in duplicates) under similar growth conditions was carried out in MM supplemented with either Glc or commercially available structural equivalent [CM3, mannotriose (Megazyme, Ireland)] of DP3 CM-β-MOS (Manβ-4Manβ-4Man deduced from ^1^H and ^13^C NMR Prabha *et al*. 2022 under review) as the sole carbohydrate source. After incubation, the culture was pelleted and washed as described in section 2.4. The washed pellet obtained was re-suspended in 1.5 ml of RNA later and stored at −80°C until further usage. RNA was extracted by using Trizol reagent + Rnaeasy mini kit as described in section 2.4. 1 μg of extracted RNA from each sample was used for the cDNA synthesis (cDNA synthesis kit, Qiagen, Germany) according to the suppliers’ protocol. Expression levels of selected differentially expressed genes in CM3 fed *B. adolescentis* were compared with Glc in thermal cycler (CFX-96, Bio-Rad) while, the 16S rRNA coding gene was used as the housekeeping control. The log_2_ fold induction level of the selected genes from both transcriptome and q-RT-PCR data was compared by running regression analysis in Excel (Microsoft).

### 2.6 Molecular docking and bioinformatics analysis

The 3D structure of the extracellular solute binding protein (bglE; BAD_RS06835) containing 432 amino acids was built using I-TASSER (Iterative Threading ASSEmbly Refinement) (Zhang 2008). 3D structures of mannobiose (M2) and mannotriose (M3) were built using carbohydrate builder module from glycam web server (Sanner 1999). Autodock 4.2 was used to perform the docking of BAD_RS06835 with M2 and M3 (Goodsell *et al*. 1996; Kirschner *et al*. 2008). Auto dock tools were used for assigning Kollman charges and adding active torsions to the protein and ligands. Water molecules were removed and polar hydrogen groups were added. No solvents were considered for docking and the protein structure was set to be rigid. The grid size was set to 92, 40, and 106 A° along the three axes (x, y, z) with the spacing of 0.375 A°. With the initial population size of 150, Lamarckian Genetic Algorithm (LGA) was employed for 100 runs in a maximum of 25, 00,000 energy evaluations. All other parameters were fixed at default settings, and the most plausible conformation of each docking was selected on the basis of estimated binding energy. The docked structures were further visualized using UCSF Chimera (Pettersen *et al*. 2004). The interacting amino acids were identified using ligplot^+^ 1.4.5 (Laskowski and Swinells 2011). Homologues of BAD_RS06835 (sharing amino acid sequence identities >55% and sequence coverage >95%) were retrieved using BLAST search and aligned computationally using CLUSTALW algorithm to build a phylogenetic tree. The sequence conservation logo of amino acids forming H-bonds in bglE was made using WebLogo, NCBI.

## 3. Results

### 3.1. *In-vitro* fermentation by *B. adolescentis* DSM 20083

In order to assess the carbohydrate preference of *B. adolescentis* DSMZ 20083 (other catalogue numbers: ATCC15703, and NCTC 11814) (Reuter 1971), *in vitro* fermentation assays were carried out on different polymeric dietary β-mannans and commercial β-manno-oligosaccharides. *B. adolescentis* DSMZ 20083 did not show any growth on all the tested polymeric dietary β-mannans when compared to the carbon-free control (Supplementary Table 2). *B. adolescentis* DSMZ 20083 grew well on CM3 (A_600nm_ 0.34) but CM2 and CM4 (up to 0.17) also supported the growth of *B. adolescentis* DSMZ 20083 when compared to carbon-free control (Supplementary Table 2).

Based on this initial assessment, we performed *in-vitro* fermentation studies with β-manno-oligosaccharides generated from polymeric dietary β-mannans viz., GMOS, GG-β-MOS CM-β-MOS, and KGM-β-MOS as described in our previous studies [Srivastava *et al*. 2017; Mary *et al*. 2019; Prabha *et al*. 2022 (under review)]. Considerable growth of *B. adolescentis* DSMZ 20083 was observed on all the other tested dietary β-manno-oligosaccharides (A_600nm_ ≥ 0.4). However, maximum growth was obtained in SDM supplemented with CM-β-MOS (A_600nm_ 0.8; log CFU ml^−1^ 8.5; final pH 5.5) (Table 1).

**Table 1.** *In vitro* fermentation of dietary β-manno-oligosaccharides by *B. adolescentis* DSMZ 20083.

| β-manno-oligosaccharide*<br>(0.1 % w/v) | Optical density<br>(A<sub>600nm</sub>) | Log CFU/ml | Final pH<sup>#</sup> |
| --- | --- | --- | --- |
| KGM-β-MOS | 0.42 ± 0.04 | 8.3 | 5 ± 0.001 |
| GG-β-MOS | 0.41 ± 0.02 | 7.4 | 5.1 ± 0.004 |
| GMOS | 0.2 ± 0.008 | 6.3 | 5.7 ± 0.002 |
| CM-β-MOS | 0.8 ± 0.06 | 8.5 | 5.5 ± 0.004 |

\* Crude short-chain β-manno-oligosaccharides were derived after hydrolysing respective β-mannans by ManB-1601 (55 °C, pH 6 under shaking conditions) followed by ethanol precipitation to remove higher DP oligosaccharides. <sup>#</sup>Final pH was estimated in the fermentation broth after 72 h of incubation.

The utilization pattern of *B. adolescentis* DSMZ 20083 was studied using structurally characterized β-manno-oligosaccharides [GMOS (Srivastava *et al*. 2017), GG-β-MOS (Mary et *al*. 2019) and CM-β-MOS Prabha *et al*. 2022 (under review); schematic structures provided in Supplementary Fig. 1)] and monosaccharides (Glc, Gal, and Man) by analysing their depletion from the growth medium using TLC. Glc was most rapidly utilized and was not found in the fermentation media after 24 h of growth whereas Gal was present till 24 h but completely disappeared at 48 h. However, Man remained un-utilized till the end of the fermentation period (72 h) (Fig. 1A). In the case of GMOS and GG-β-MOS, we could see a reduction in the concentration of both DP2 and DP3 species between 24-72 h of fermentation. However, a very distinctive utilization of DP2 and DP3 CM-β-MOS from the fermentation media was seen after 24 h of incubation till 72 h (Fig. 1B). Spot intensities of DP 4-6 CM-β-MOS were also seen to become lighter with the increase in fermentation time (Fig. 1B). The lighter spot intensities obtained in GG-β-MOS depletion were due to the supplementation by low initial substrate concentration (0.1 % w/v). Since the utilization of CM-β-MOS was found better than other tested dietary β-manno-oligosaccharides, we proceeded further with a growth curve analysis of *B. adolescentis* DSMZ 20083 on CM-β-MOS, control sugars (Glc and Gal), and carbon-free MM wherein, *B. adolescentis* DSMZ 20083 showed preferential utilization and rapid growth on CM-β-MOS when compared to Gal. However, the onset of exponential phase on CM-β-MOS was delayed when compared to Glc. Carbon-free MM was unable to sustain the growth of *B. adolescentis* above A_600nm_ ~0.2. The generation time of *B. adolescentis* DSMZ 20083 on CM-β-MOS was found to be 220 min whereas on Gal and Glc it was 380 min and 60 min, respectively (Fig. 2A & B).

**Figure 1.**
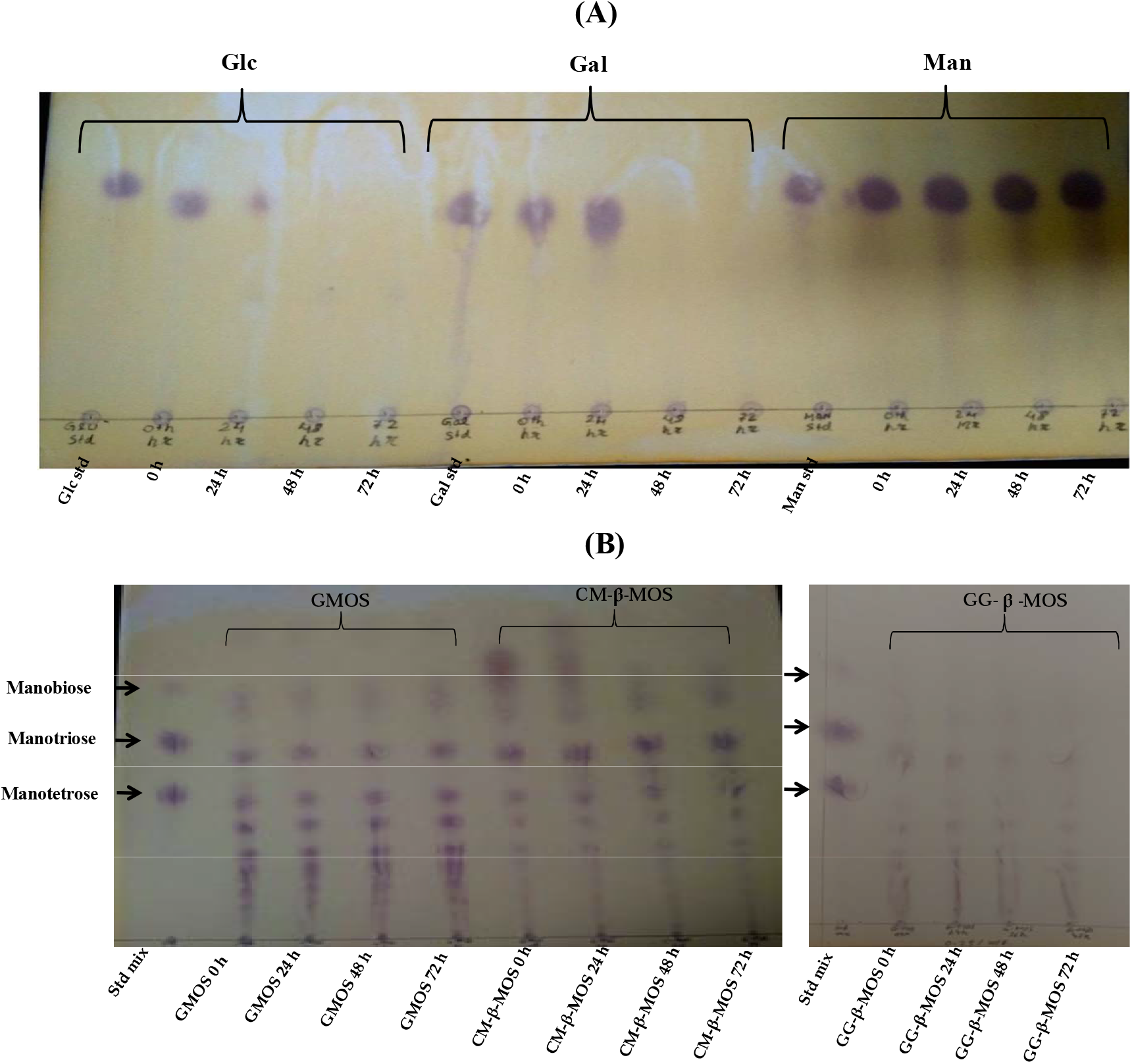
(A) Depletion patterns of supplemented monosaccharides (Glc, Gal, and Man) from MM by *B. adolescentis* DSMZ 20083 up to 72 h of growth. Lanes 1, 6, and 11 represent Glc, Gal, and Man standards, respectively. Lanes 2 to 5, 7 to 10 and 12 to 15 represent the samples harvested from Glc, Gal and Man supplemented MM, respectively at defined time intervals. (B) Depletion patterns of supplemented dietary β-mannooligosaccharides from MM by *B. adolescentis* DSMZ 20083 up to 72 h of growth. Lanes 1 and 10 represent detection standard comprising of mannobiose, mannotriose, and mannotetrose. Lanes 2 to 5, 6 to 9, and 11 to 14 represent GMOS, CM-β-MOS, and GG-β-MOS, respectively. Schematic structures of β-mannooligosaccharides are provided in Supplementary Figure 1.

**Figure 2.**
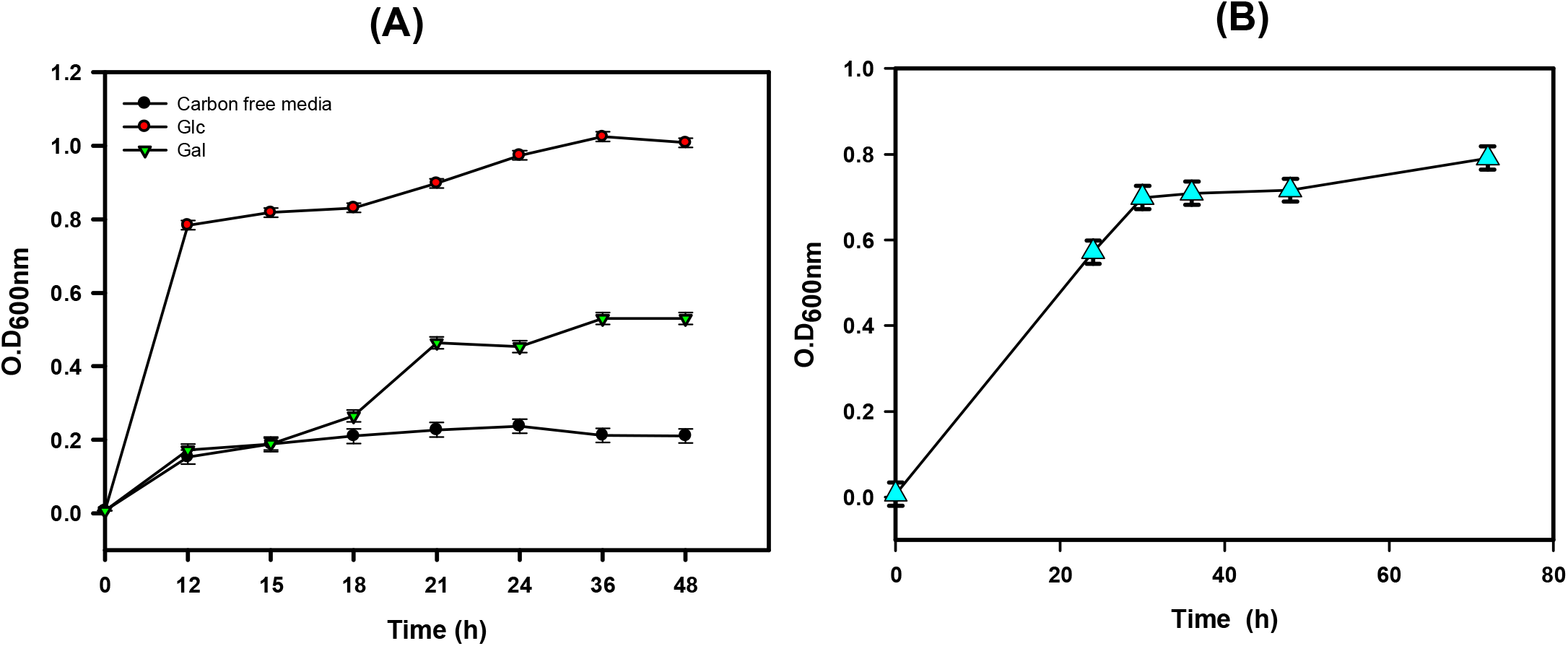
(A) Growth curves of *B. adolescentis* DSMZ 20083 on Glc, Gal, and carbon-free media. (B) Growth curve of *B. adolescentis* DSMZ 20083 on CM-β-MOS.

### 3.2. Microarray and q-RT-PCR analysis

To decipher the underlying molecular mechanism by which *B. adolescentis* DSMZ 20083 utilizes the un-decorated, short-chain CM-β-MOS (majorly DP2, DP3, and DP4); a whole-genome transcriptional analysis was carried out. Volcano plot analysis showed that growth on CM-β-MOS induced a significant (*P* < 0.05) differential gene regulation versus Glc wherein a total of 632 differentially expressed genes (DEGs) were identified (Fig. 3A). Heat map analysis of these DEG’s illustrated that 164 genes were up-regulated (fold change ≥ 2) while, 85 genes were found down-regulated (fold change ≤ −2) (Supplementary Fig. 2). The larger number of DEGs pointed towards the global effect of CM-β-MOS on the overall metabolism of *B. adolescentis* DSMZ 20083 (Lei *et al*. 2018). The top fifteen up-regulated genes in the transcriptome analysis have been provided in Table 2. Among the up-regulated genes, we were able to assign three distinct gene clusters (Fig. 3B & 4A); cluster I [Three genes: BAD_RS06080, cscB (BAD_RS06085), cscA (BAD_RS06090)], cluster II [Five genes: bglR (BAD_RS06840), bglE (BAD_RS06835), bglF (BAD_RS06830), bglG (BAD_RS06825), and bglB (BAD_RS06820)] and cluster III (Seven genes: bfrB (BAD_RS07050), bfrC (BAD_RS07045), bfrD (BAD_RS07040), PF04854 (BAD_RS07035), bfrR (BAD_RS07030), bfrA (BAD_RS07025)] which are possibly employed by *B. adolescentis* DSMZ 20083 for the uptake and deconstruction of CM-β-MOS into monosaccharides. Referring to the annotations provided in KEGG, CAZy, and NCBI databases, these operons code for proteins such as Lacl family transcriptional regulator (BAD_RS06840 and RS06080), Major facilitator superfamily (MFS) transporter (BAD_RS06085), ATP-binding cassette (ABC) transport system comprising of extracellular solute binding protein (BAD_RS06835 & BAD_RS07050) and permease domains [multi-species ABC transporter permeases (BAD_RS06830 & BAD_RS07045), & multi-species carbohydrate transporter permeases (BAD_RS06825 & BAD_RS07040)], and glycoside hydrolases belonging to GH1 β-glucosidase (BAD_RS06820) and GH32 β-fructofuranosidase (BAD_RS06090 & BAD_RS07025). According to RegPricise3.0, clusters I and II (highest up-regulated in presence of CM-β-MOS) are mapped to be involved in sucrose and β-glucosides utilization pathways, respectively whereas, cluster III has been previously mentioned to be involved in the uptake and catabolism of fructo-oligosaccharides (Sela *et al*. 2008). The expressed glycoside hydrolases upon analysis using SignalP 5.0 were found not to have any signal peptides and thus would occur most likely intra-cellularly. Surprisingly, the possible cell-attached GH26 endo-β-mannanase predicted to be involved in β-mannan catabolism (Kulcinskaja *et al*. 2013) was found down-regulated (−0.669 times) in comparison to the gene expression observed in the presence of Glc.

**Figure 3.**
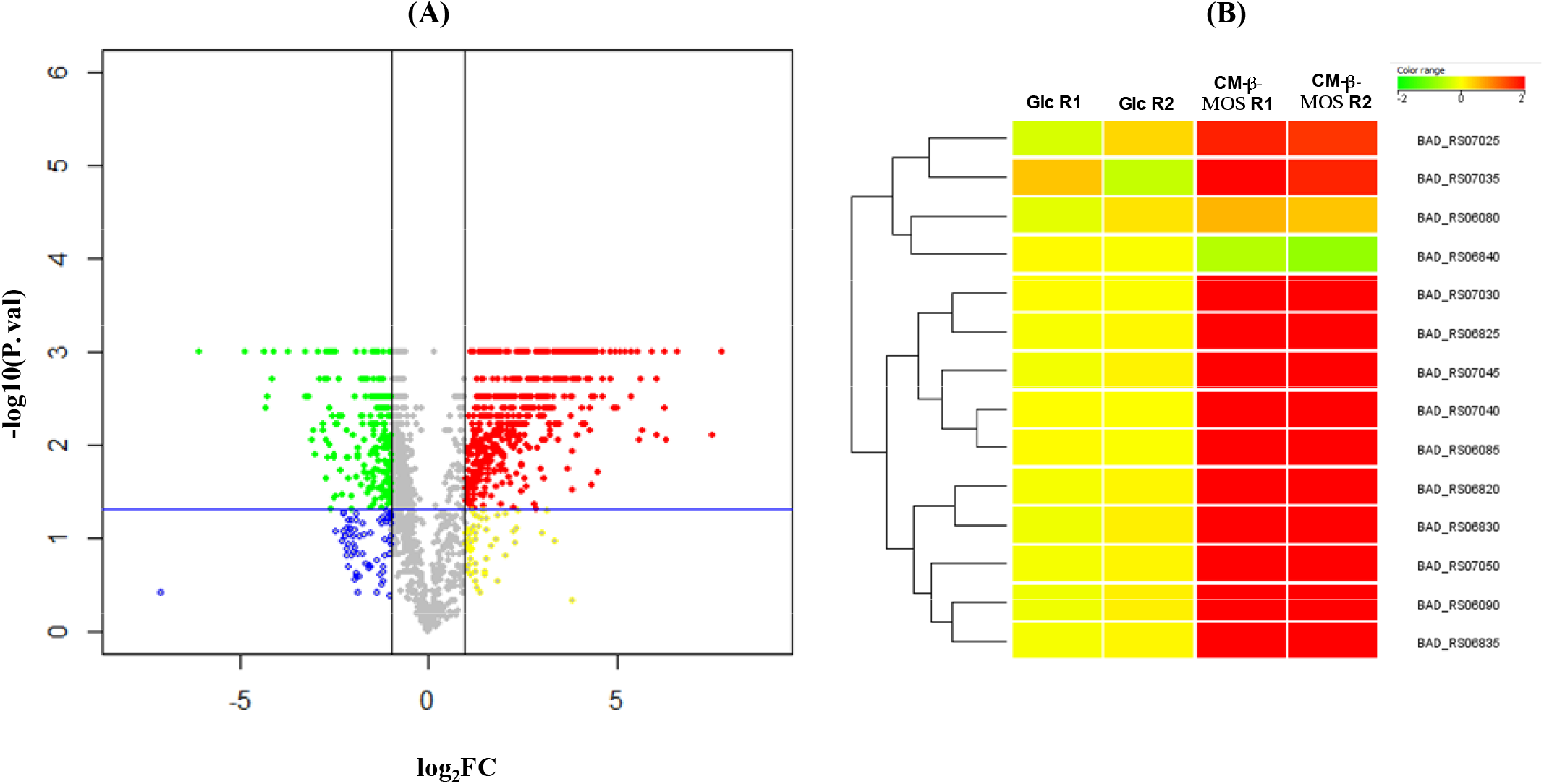
(A) Volcano plot for differential gene expression in *B. adolescentis* grown on CM-β-MOS versus Glc. The x axis indicates the differential expression profiles by plotting the fold-change at log_2_ scale. The y axis indicates the statistical significance for the differences obtained in the gene expression (P value from a *t*-test) on log_10_ scale. Each dot in the plot represents a single gene. Blue: Fold change ≤ −1 and *P* > 0.05; Yellow: Fold change ≥1 and *P* > 0.05; Green: Fold change ≤ −1 and *P* < 0.05 and Red: Fold change ≥1 and *P* < 0.05. (B) Heat map comparison of highly up-regulated gene clusters involved in the uptake and catabolism of CM-β-MOS versus Glc in *B. adolescentis* DSMZ 20083. R-replicate.

**Table 2.** Top 15 significantly up-regulated genes in *B. adolescentis* DSMZ 20083 during growth on CM-β-MOS when compared to Glc as assessed by the whole genome transcriptome analysis using microarray.

| S.No. | Locus ID | Log<sub>2</sub> fold change* | P value | Annotation** |
| --- | --- | --- | --- | --- |
| 1 | BAD_RS05300 | 7.8 | 0.00 | Formate C-acetyltransferase |
| 2 | BAD_RS05460 | 7.7 | 0.00 | MULTISPECIES: helix-turn-helix transcriptional regulator |
| 3 | BAD_RS05115 | 7.3 | 0.00 | Multispecies ROK family protein |
| 4 | BAD_RS06820 | 7.3 | 0.00 | GH1 β-glucosidase |
| 5 | BAD_RS06830 | 6.9 | 0.001 | Multispecies: sugar ABC transporter permease |
| 6 | BAD_RS08375 | 6.9 | 0.01 | MULTISPECIES: sn-glycerol-3-phosphate ABC transporter ATP-binding protein |
| 7 | BAD_RS03815 | 6.7 | 0.01 | Glucose-1P-adenyltransferase |
| 8 | BAD_RS00260 | 6.4 | 0.01 | Hsp20 alpha crystalline family protein |
| 9 | BAD_RS01100 | 6.3 | 0.01 | Multispecies: 30S ribosomal protein S16 |
| 10 | BAD_RS07760 | 6.3 | 0.01 | Glutamyl-Q tRNA (Asp) synthetase |
| 11 | BAD_RS05395 | 6.1 | 0.01 | RNA polymerase sigma factor |
| 12 | BAD_RS06835 | 6.08 | 0.00 | Extra cellular solute binding protein |
| 13 | BAD_RS05920 | 5.9 | 0.001 | Lactate dehydrogenase |
| 14 | BAD_RS03185 | 5.9 | 0.01 | Cysteine synthase |
| 15 | BAD_RS04160 | 5.8 | 0.01 | Peptide deformylase |

\*Significance of fold change data is judged by having a P value of no more than 0.01. \*\*Gene annotations were ascertained by blast analysis using Swiss prot and are listed by the order of magnitude of induction attained.

**Figure 4.**
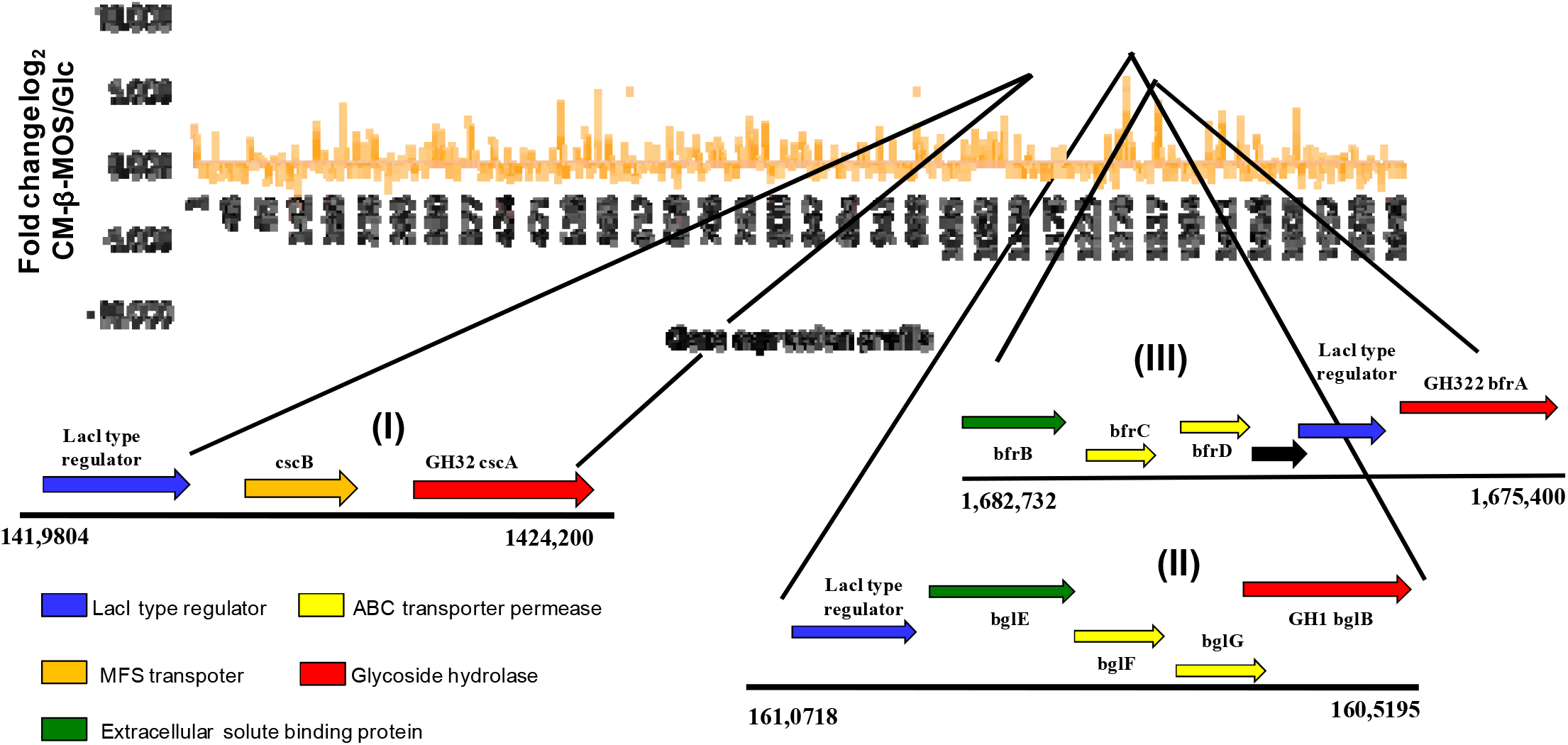
Upper panel: Whole genome transcriptome map of differentially expressed genes after growth on CM-β-MOS versus Glc in *B. adolescentis* DSMZ 20083. Lower panel: Gene organization of assigned clusters involved in the uptake and catabolism of *B. adolescentis* DSMZ 20083.

Genes involved in the catabolism of galactose via the Leloir pathway (Fortina *et al*. 2003) such as aldose-1-epimerase (BAD_RS05750), galactokinase (BAD_RS07055), UDP-glucose-4epimease (BAD_RS0391), and phosphoglucomutase (BAD_RS01935) were also found up-regulated (*P* < 0.05, fold change ≥ 3). Repressor Open Reading Frame Kinase (ROK) family protein (BAD_RS05115), which is known to act either as transcriptional activators or kinases, was the second most highly up-regulated gene (7.3 fold) (O’Conell *et al*. 2014). According to Gene ontology analysis, 185, 175 and 174 participating genes involved in various biological, cellular, and metabolic processes, respectively among others were found up-regulated (Supplementary Fig. 3A-C Up-regulated; D&E Down-regulated). KEGG enrichment analysis highlighted the up-regulation of important metabolic pathways involved in the metabolization of carbon source viz., biosynthesis of secondary metabolites; carbon metabolism; galactose metabolism; ABC transport system; pyruvate metabolism; starch and sucrose metabolism; glycolysis/gluconeogenesis; pentose and glucouronate interconversions among others (Supplementary Fig. 4A). Pathways related to the biosynthesis of amino acids, vitamin B6 metabolism, and homologous recombination, and few ABC transport systems were found to be down-regulated in comparison to the Glc control (Supplementary Fig. 4B).

To corroborate whether *B. adolescentis* DSMZ 20083 shows a preference for DP3 as seen with commercial β-manno-oligosaccharide (Supplementary Table 2), we fed purified DP2 and DP3 β-MOS generated from guar gum (Mary *et al*. 2019) and DP4 β-MOS from locust bean gum (Srivastava *et al*. 2017) to *B. adolescentis* DSMZ 20083 (experimental details provided as a foot note in Supplementary Table 2) and found higher growth on DP3 when compared with DP2 and DP4 β-MOS (Supplementary Table 2). This made us curious to know whether DP3 is the most bifidogenic β-MOS and can induce the uptake and metabolic pathways for the utilization of β-MOS as found in microarray analysis, we performed q-RT-PCR studies of *B. adolescentis* DSMZ 20083 passaged on CM3 which is a structural equivalent of DP3 CM-β-MOS. As anticipated, all selected nine genes related to the uptake and metabolism of CM-β-MOS showed significant up-regulation (*P* < 0.05) in the q-RT-PCR studies when compared to Glc (Fig 5). MFS transporter cscB showed higher up-regulation (6.9 fold) in comparison to bglE (3.4 fold) pointing that MFS could be the primary transporter of DP3 β-manno-oligosaccharide in *B. adolescentis* DSMZ 20083 as previously in *Bifidobacterium lactis* Bl-04, MFS transporter was found up-regulated 17 folds higher upon growth on galacto-oligosaccharides having DP ≥ 3 (more than 94%) when compared to gentiobiose (DP2) (Anderson *et al*. 2013). The cell-attached GH26 endo-β-mannanase (BAD_RS05480) was analysed again to confirm its possible expression however, it did not get induced (Cq value similar to no template control) in response to growth on CM-β-MOS and corroborated well with the microarray data. The coefficient of determination value (R^2^ = 0.812) suggested that up-regulation observed for the selected genes in the microarrays and q-RT-PCR analysis had a good correlation.

**Figure 5.**
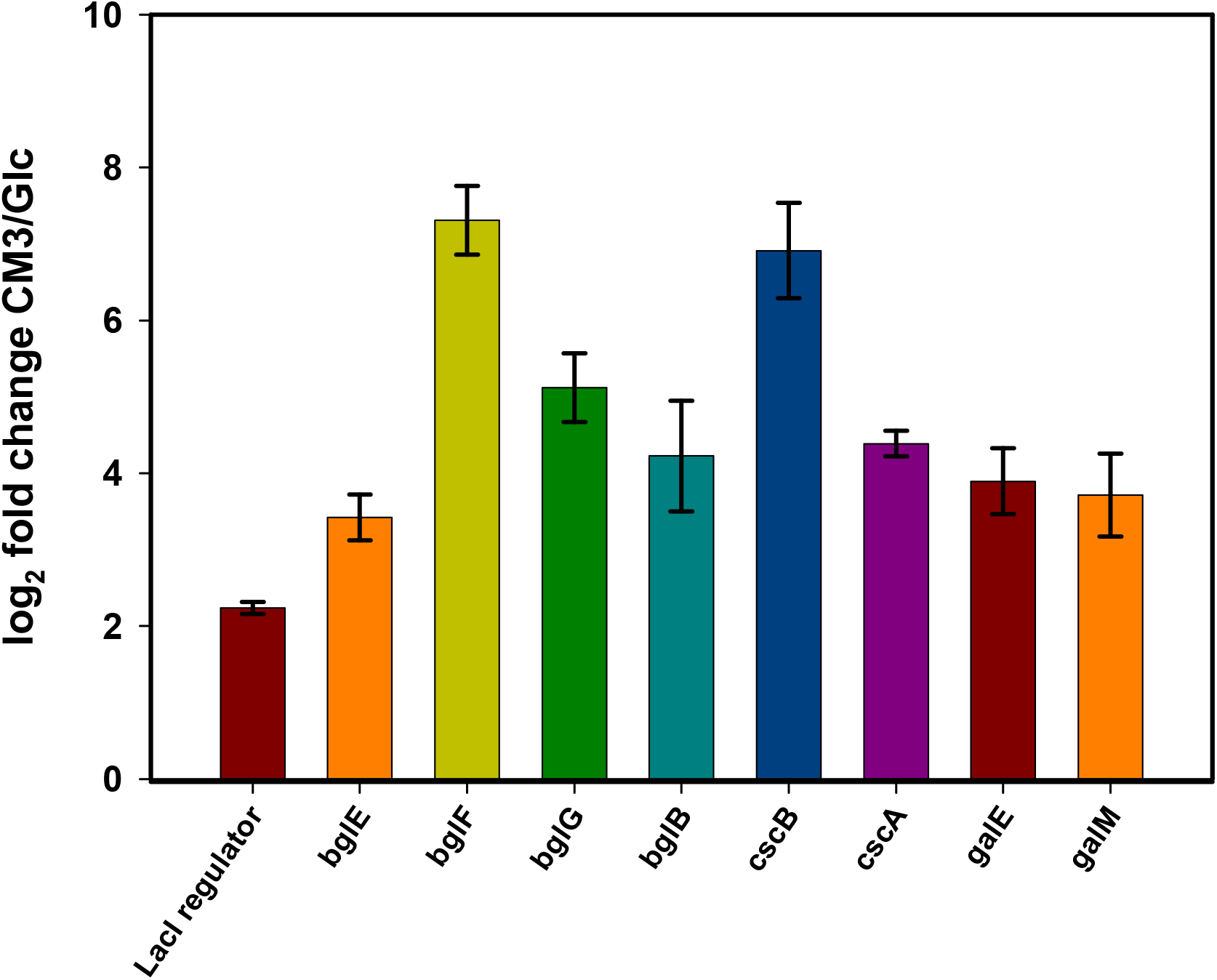
Log_2_ fold change of selected differentially up-regulated genes involved in the utilization of CM3 by *B. adolescentis* DSMZ 20083 using q-RT-PCR analyses.

### 3.3 Molecular docking and bioinformatics analysis

The binding interactions for ESBP (BAD_RS06835) with β-manno-oligosaccharides (M2 and M3) were investigated by computational docking studies. M2 displayed a low free binding energy of −1.73 kcal/mol suggesting strong electrostatic interactions with ESBP. Hydrogen bonds played an important role in binding of the ligand to receptor. M2 interacted with hydrophobic residues [Trp393-2.81 Å, Ala394-2.56 Å] polar residues [Ser397-2.78 Å, Thr111-2.56 Å, and Ser398 (2 hydrogen bonds: 2.93 Å) and non-polar residue (Trp112-2.92 Å) of ESBP by hydrogen bonding interaction. The binding pocket (4Å) accommodating M2 was surrounded by thr204, leu208, phe207, trp393, ala394, trp112, trp86, and his90 (Fig. 6A & B). M3 showed higher binding energy (−0.23 kcal/mol) and weaker electrostatic interaction when compared to M2. It formed hydrogen bonds with Ser87 (4 hydrogen bonds; 2.85Å, 2.71Å, 2.85Å, and 2.71Å), Trp86 (2 hydrogen bonds: 2.96 Å and 2.96 Å) Trp112 (2 hydrogen bonds; 3.19 Å, and 3.19 Å), Thr111 (2 hydrogen bonds; Å, 2.83 and 2.83 Å) and Gly318 (4 hydrogen bonds; 2.63 Å, 2.78 Å, 2.63 Å, and 2.78 Å) during interaction with ESBP. The binding site was found to be surrounded by Gly109, Thr110, and Phe315 (Fig 6C & D). In contrast, ABC transporter, *Bl*MnBP2 from *B. animalis subsp. lactis* showed higher binding affinity towards mannotriose as compared to mannobiose (Ejby *et al*. 2019).

**Figure 6.**
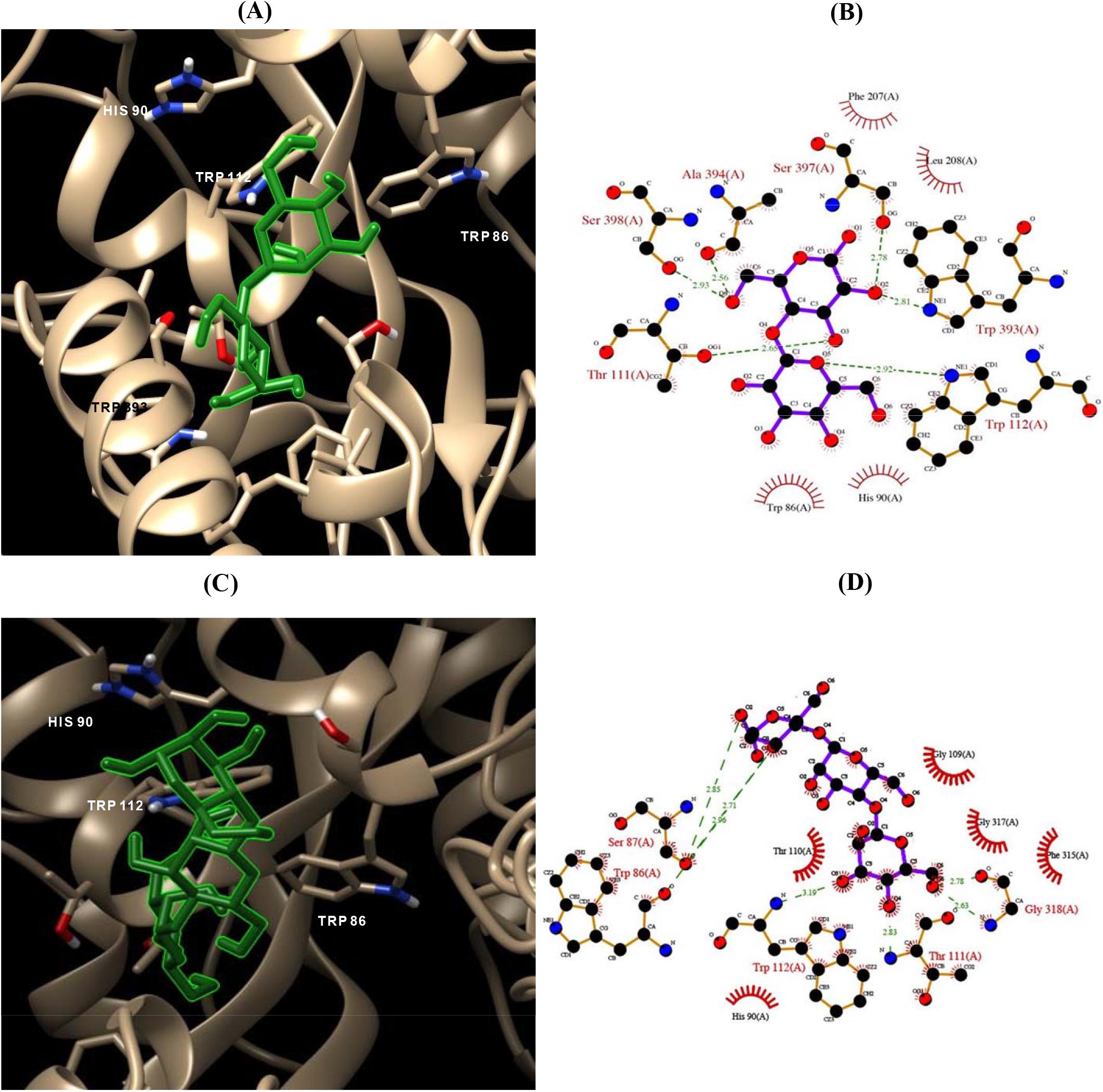
Molecular docking studies of mannobiose and mannotriose with bglE to estimate the binding energy, hydrogen bonds and neighbouring amino acid residues. Closeup of binding pocket residues involved in the interaction of bglE with (A) M2 and (B) M3. LIGPLOT representation of hydrophobic and hydrogen bonding interactions between the amino acid residues at carbohydrate binding site of bglE with (C) M2 and (D) M3. Red arcs indicate hydrophobic interactions while, H-bond interactions are given by the green dashed lines.

BAD_RS06835 from *B. adolescentis* ATCC 15703 was found closely related to other ESBP’s homologues from *Bifidobacterium pseudocatenulatum, Bifidobacterium lemurum, Bifidobacterium pseudolongum*, and *Bifidobacterium angulatum* by sharing a high sequence similarity of 94.91%, 92.11% and 91.20 %, respectively. However, ESBP from *Bifidobacterium asteroids, Bifidobacterium longum*, and *Bifidobacterium breve* were found distantly related to BAD_RS06835 (Supplementary Figure 5A). A high degree of conservation was seen in bglE homologues of analyzed *Bifidobacterium* sp. for amino acid present at position 111 (threonine) and 393 (alanine). Amino acids at positions 112, 394, 397, and 398 were also found fairly conserved within most bifidobacteria. The conserved amino acids present in the ESBP’s from other *Bifidobacterium* sp. can also potentially form hydrogen bonds with mannobiose and help in its capturing (Supplementary Figure 5B & C).

## 4. Discussion

*Bifidobacteria* spp. are key gut commensals and recognized as beneficial microbes by the virtue of their pivotal contributions made towards gut microbiota homeostasis both pre- and post-weaning (Turroni *et al*. 2018). Gaining functional insights into mannanolytic abilities of these beneficial bacteria can provide a strong foundation for using β-mannan / β-manno-oligosaccharide based dietary interventions to modulate the adult gut microbiome. Among the bifidobacterial species, probiotic *B. animalis* subsp. *lactis* has been reported to be mannanolytic and can utilize carob galactomannan (Ejby *et al*. 2019). Recently, it has been demonstrated that *B. adolescentis* DSMZ 20083 is able to utilize short-chain oligosaccharides derived from β-mannan sources like acetylated spruce wood, cassia gum, and low viscosity LBG (La Rosa *et al*. 2019; Li *et al*. 2020; & Bhattacharya *et al*. 2021). However, to date a molecular understanding of β-mannano-oligosaccharide utilization in adult-associated *B. adolescentis* DSMZ 20083 is lacking.

In the present study initially, the ability of *B. adolelscentis* DSMZ 20083 to assimilate β-manno-oligosaccharide generated from common dietary β-mannans such as LBG, guar gum, konjac, and copra meal was investigated. All the β-manno-oligosaccharides; GMOS, GG-β-MOS, KGM-β-MOS, and CM-β-MOS supported the growth of *B. adolescentis* DSMZ 20083 appreciably but maximum growth was attained on media supplemented with CM-β-MOS (Table 1). In TLC-based analysis of oligosaccharide depletion from the growth medium, *B. adolescentis* DSMZ 20083 was found to distinctly utilize both DP2 and DP3 CM-β-MOS. It also suggested perceptible but poor utilization of higher DP oligosaccharides (≥ DP4). This pattern of utilization of un-substituted CM-β-MOS in *B. adolescentis* DSMZ 20083 was found distinctive from the substituted GMOS or GG-β-MOS where higher oligosaccharides remained un-utilized (Fig 1B). However, fermentation of pure DP2, DP3, and DP4 β-MOS were indicative of preferential utilization of DP3 over DP2 and DP4 (Supplementary Table 2). Data obtained from oligosaccharide depletion studies and *in vitro* fermentation aligned well with the previous reports which showed that *B. adolescentis* DSMZ 20083 is a short-chain oligosaccharide forager (Mei *et al*. 2011). TLC analysis of monosaccharide utilization by *B. adolescentis* DSMZ 20083 showed expectedly good utilization of Glc and Gal and at the same time no growth on Man (Fig. 1A) as reported previously (Turroni *et al*. 2012). This was intriguing as mannose forms the building block of β-manno-oligosaccharides which the bacterium was found to utilize well. Previously, *Bifidobacterium lactis* BB12 has also been reported to grow well on oligomers (xylo-oligosaccharides) but failed to utilize the monomer (xylose) (Crittenden *et al*. 2002). Certain members of *Bifidobacteria* sp. are not able to grow on some monosaccharides due to the lack of specific transporters associated with their import (Turroni *et al*. 2012; Zeybek *et al*. 2020) but still can internalize and metabolize oligosaccharides composed of such monomers by virtue of specialized genetic loci comprising of high-affinity specific transporters and glycoside hydrolases (Gilad *et al*. 2010).

Single colour microarray analysis allowed us to unravel the global gene expression patterns and reconstruct the uptake and catabolic pathway utilized by *B. adolescentis* DSMZ 20083 to grow on CM-β-MOS (Fig. 7). Growth on linear β-1,4 mannosyl linked CM-β-MOS by *B. adolescentis* DSMZ 20083 led to the up-regulation of three distinct gene clusters coding for proteins such as Lacl family transcriptional regulator, MFS transporter, ATP-binding cassette transport system comprising of extracellular solute binding protein with associated permease domains and glycoside hydrolases belonging to GH1 β-glucosidase and GH32 β-fructofuranosidases (Fig 3B & 4A). Up-regulation or induction of such genes is typical to the carbohydrate utilization machinery expressed by *Bifidobacteria* sp. in response to glycans (Fushinobu and Haschem, 2021). Up-regulation of two different genetic loci with similar architecture in response to growth on galacto-oligosaccharide having a varied degree of polymerization (DP 2-8) have been reported previously in *Bifidobacterium lactis* Bl-04 and *Bifidobacterium breve* (Anderson *et al*. 2013; O’Connel *et al*. 2013). However, the strategy adopted by *L. plantarum* WCFS1 which utilized only a single locus for internalizing unsubstituted GMOS (Panwar and Kapoor 2019) was in contrast.

**Figure 7.**
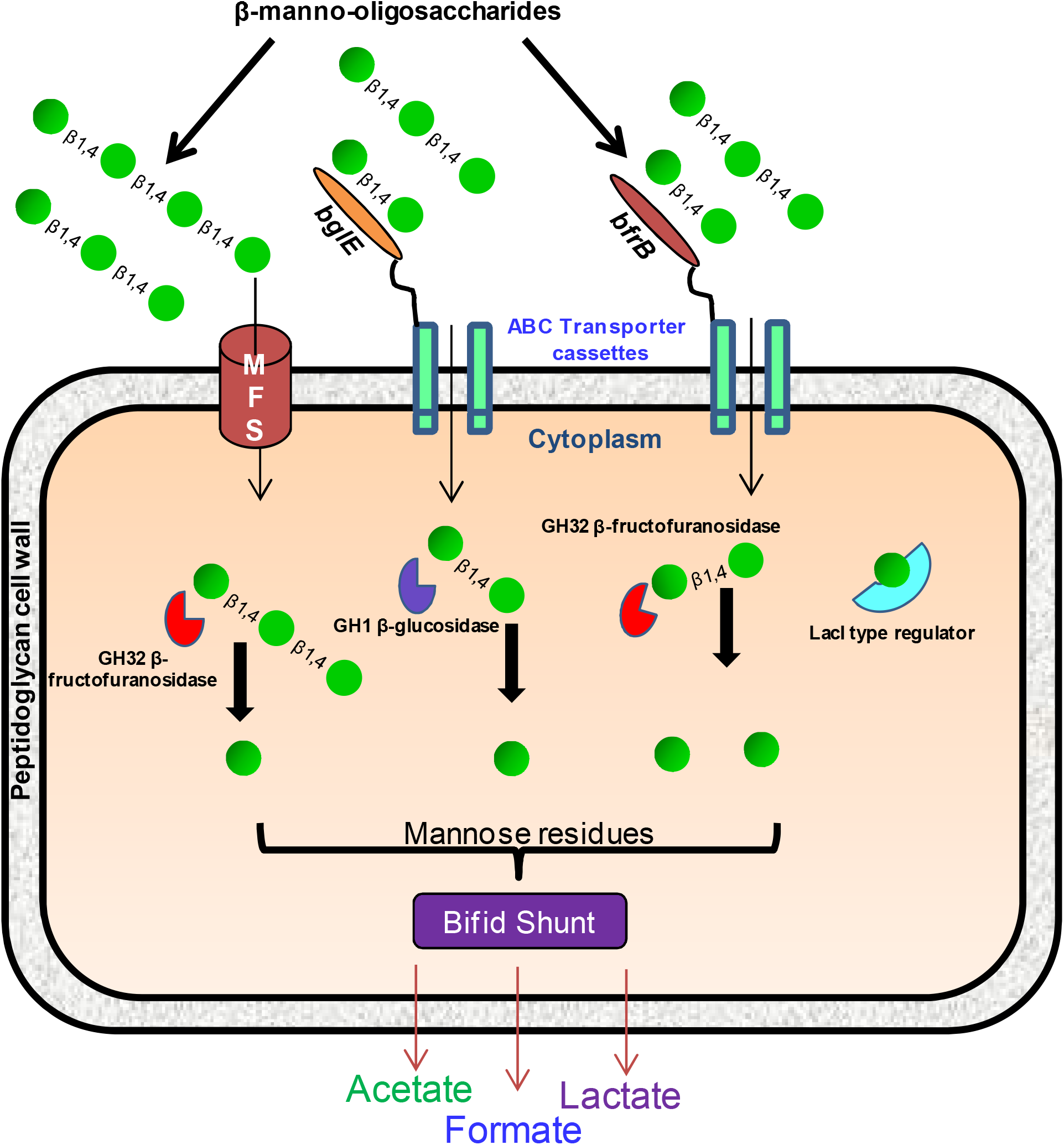
Proposed schematic model for the uptake and deconstruction of short chain CM-β-MOS in *B. adolescentis* DSMZ 20083. Extracellular solute binding protein, bglE located in cluster II possibly captures DP2 majorly and to some extent DP3 which are then transported by the associated permeases. bfrB in cluster III could also function in a similar manner. MFS transporter located in cluster I would transport β-manno-oligosaccharides having DP ≥ 3. Once inside the cytoplasm, the oligosaccharides could be cleaved by exo-acting enzymes GH1 β-glucosidase and GH32 β-fructofuranosidases present in cluster I, II or III. The generated D-mannose could possibly enter amino sugar/nucleotide sugar metabolism or the bifid shunt.

ABC transporter cassette comprising of a lipid anchored extracellular solute binding protein (ESBP; bglE) and two associated carbohydrate permeases (bglF and bglG) in cluster II predicted to be involved in the uptake and catabolism of glucosides was probably responsible for the uptake and transport of DP2 and to lesser extent DP3 CM-β-MOS as bglE showed higher binding affinity for DP2 than DP3 CM-β-MOS (Fig 6A & C). ESBPs from *Bifidobacterium* sp. which are characterised to date are glycan-specific and bind their cognate oligosaccharide ligands tightly (Ejby *et al*. 2016). Cluster III, a fructo-oligosaccharide utilization operon, which encodes the complete ATP binding cassette comprising of ESBP and transporter permeases also got up-regulated upon growth on CM-β-MOS which is consistent with its functional role (Sela *et al*. 2008). In Bifidobacteria, the lipid anchored ESBP extends through the thick peptide glycan layer and directly captures dietary polysaccharides and oligosaccharides (Van der meullen *et al*. 2006). The associated permeases made up of two trans-membrane domains actively port the acquired glycans into the cytoplasm coupled with ATP-hydrolysis by the two cytoplasmic nucleotide-binding domains. Specificity of these ABC transporters towards the substrate arises from its trans-membrane domains which do not share sequence homology with other transporters (Saier 2000). Transporters targeting β-mannooligosaccharides are relatively less prevalent among *Bifidobacterium* sp. but if present may favour metabolic specialisation within the species (Fushinobu and Hachem 2021). MFS transporters have been previously known to transport short-chain oligosaccharides in *Bifidobacterium lactis* Bl-04. Growth up on galacto-oligosaccharides (DP ≥ 3 more than 94%) up-regulated MFS transporter 17 folds higher when compared to gentiobiose (DP2) in *Bifidobacterium lactis* Bl-04 (Anderson *et al*. 2013). This suggested that MFS preferably transport higher DP oligosaccharides. From this information and q-RT-PCR studies, we inferred that internalization of CM-β-MOS (DP ≥ 3) is likely to be carried out primarily via the MFS transporter located in cluster I and thus makes it a novel route for the uptake of β-manno-oligosaccharides, as till date, only PTS and ABC systems have only been reported to be involved in the uptake of β-manno-oligosaccharides (Ejby *et al*. 2019; Panwar and Kapoor, 2019).

Glycoside hydrolases (GH) which can target the glycans and release monosaccharide products are encoded alongside ABC and MFS-transporters (van den Broek *et al*. 2008; Ejby *et al*. 2013). Once inside the intracellular space, DP2 and DP3 CM-β-MOS are possibly hydrolysed to mannose residues by the GH1 β-glucosidase present in cluster II or GH32 β-fructofuranosidases present in both cluster I & III. GH1 β-glucosidases are exo-acting enzymes previously shown to cleave mannose residues from the non-reducing end in β-mannans (Bauer *et al*. 1996). However, GH1 β-glucosidase though present in the β-mannan utilization cluster I of *B. animalis* sub sp. *lactis* was not found to be up-regulated upon growth on galacto- and glucomannans, possibly due to the availability of a functional mannosidase and a surface-attached GH5 endo-β-mannase which can contribute towards the deconstruction of β-mannan. The β-glucosidase from *B. brevis* UCC2003 has been shown to be active on β-1,4 linked cellobiose and cellodextrin (β-1,4-glucooligosaccharides) (Pokusaeva *et al*. 2011). Exo-acting GH32 β-fructofuranosidase (BAD_RS06090) was found to be up-regulated previously in *B. adolescentis* DSMZ 20083 upon growth on xylo-oligosaccharides suggesting its potential role in cleaving β-linked oligosaccharides (Yang *et al*. 2019). In *Bifidobacteria*, there is more prevalence of exo-acting enzymes capable of releasing monosaccharide’s from the non-reducing end of oligosaccharides although, a few endo-acting enzymes have also been reported (van den Broek *et al*. 2008). Since *B. adolescentis* DSMZ 20083 utilizes only short-chain oligosaccharides; the up-regulation of exo-acting enzymes in the present study seems justified. The previously reported GH26 endo-β-mannanase (Kulcinskaja *et al*. 2013) was not found to be up-regulated both in the transcriptome and q-PCR analysis but was found up-regulated after growth on xylo-oligosaccharide (Yang *et al*. 2019) thus making its functionality unclear. The utilization pattern of the previously known β-mannan utilizer *B. animalis* subsp. *lactis* appears inherently different from *B. adolescentis* DSMZ 20083 as the former can degrade polymeric β-mannan extracellularly and internalize the generated oligosaccharides (Ejby *et al*. 2019) while, the latter was found capable of growing only on β-manno-oligosaccharides of short-chain length (DP2 and DP3 CM-β-MOS). Here, the pattern of utilization draws a parallel with that of DP2 GMOS derived from locust bean gum in *L. plantarum* WCFS-1 in which a GH1 6-P-β-glucosidase is predicted for the cleavage of DP2 GMOS. However, the oligosaccharides were imported directly presumably using a PTS transport system (Panwar and Kapoor 2019). Such skewed utilization patterns in *Bifidobacteria* sp. and *Lactobacillus* sp. can be interpreted as a method of co-existence and partially avoiding competition with the generalists such as *Bacteroides ovatus, Roseburia intestinalis* etc. that target polymeric substrates in the competitive gut ecosystem (Ejby *et al*. 2016 and La Rosa *et al*. 2019).

As per the annotations in the KEGG pathway, the generated D-mannose from β-manno-oligosaccharides can enter amino sugar/nucleotide sugar metabolism or the bifid shunt. However, the actual route by which this is achieved still needs further probing as no known enzymes which are capable of converting mannose to glucose or fructose like cellobiose epimerase, mannose 6-P-isomerase, xylose isomerise, etc. are present in *B. adolescentis* DSMZ 20083 or if present were not found up-regulated. Our transcriptome data interestingly revealed the up-regulation of genes involved in the Leloir pathway of galactose metabolism. One protein of this pathway viz., phospho-glucomutase, which directs glucose-1-P into the glycolytic pathway, has recently been reported to also act on mannose-1-P and convert it to mannose-6-P in the β-mannan utilization pathway of the primary degrader *Roseburia intestanalis* (La Rosa *et al*. 2019). Interestingly, ROK family protein which regulates phosphoglucomutase (BAD_RS01935) (RegPrecise 3.0) was also found highly up-regulated indicating their possible role in CM-β-MOS metabolism in *B. adolescentis* DSMZ 20083 but this needs further investigation.

## Conclusions

In the present study, we employed whole-genome transcriptome analysis to identify the operons involved in the utilization of β-manno-oligosaccharide by *B. adolescentis* DSMZ 20083. We identified three gene clusters namely sucrose, β-glucoside and fructo-oligosaccharide utilization operons encoding MFS & ABC transport systems to internalise, as well as GH1 (β-glucosidase) & GH32 (β-fructofuranosidase) glycoside hydrolases to deconstruct the linear, short-chain length CM-β-MOS. The insights gained on the metabolism of dietary β-manno-oligosaccharides by adult gut associated *B. adolescentis* DSMZ 20083 can be instrumental in developing personalized nutrition for precision microbiome-based interventions, which eventually may prove useful in ameliorating human diseases manifested by the gut dysbiosis.

## Supporting information

Dupplementary Figure 4

## Acknowledgements

We thank Director, CSIR-CFTRI, for constant encouragement and support. MK acknowledges the financial support of the project (BT/PR23266/PFN/20/1286/2017; GAP No. 523) funded by the Department of Biotechnology, Ministry of Science and Technology, Govt. of India. PRM thanks UGC, Govt. of India for the award of Senior Research Fellowship (SRF). MP thanks DBT, New Delhi for the award of SRF. The authors declare that they have no known conflict of interest.

## AUTHOR’S CONTRIBUTION

P.R.M. and M.K. conceived and designed the research. P.R.M conducted all the wet lab experiments and M.P performed the Bioinformatics analyses. P.R.M, M.P, and M.K analysed the data. P.R.M. and M.K. wrote and drafted the manuscript. All authors read and approved the final version of the manuscript.

