## Supplementary Figure 1 for "Insights into β-manno-oligosaccharide uptake and metabolism in *Bifidobacterium adolescentis* DSMZ 20083 from whole-genome microarray analysis"

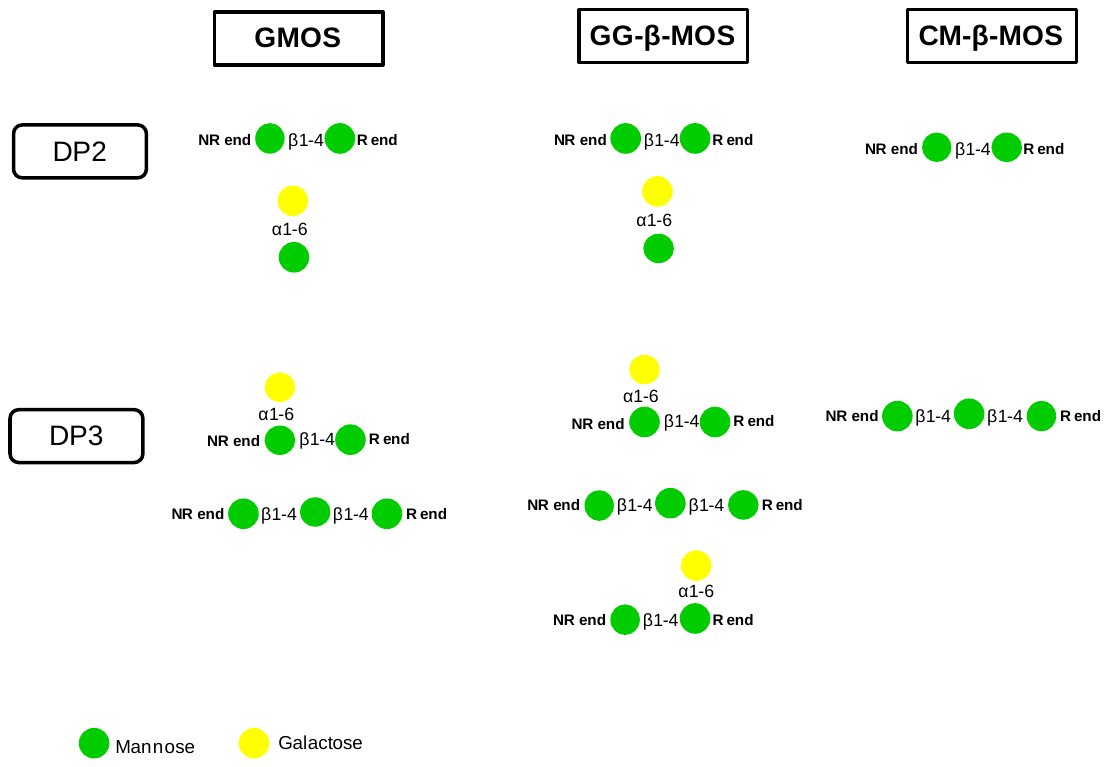

**Figure 1** Schematic structures of GMOS, CM-β-MOS and GG-β-MOS. Monosaccharides are colour coded as per symbol nomenclature for glycans (SNPG). D-Mannose; green D-Galactose; yellow. NR-non reducing end and R-reducing end.
