## Supplementary figures and images for "Insights into β-manno-oligosaccharide uptake and metabolism in *Bifidobacterium adolescentis* DSMZ 20083 from whole-genome microarray analysis"

### Supplementary Figure 3

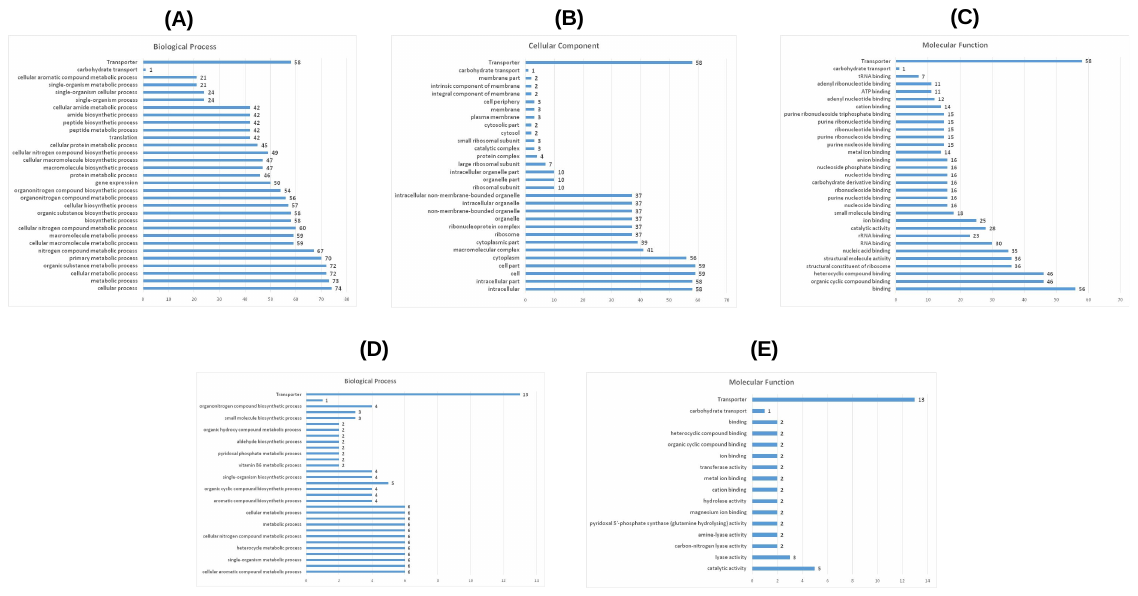
**Supplementary Figure 3**
