## Supplementary material for "Insights into β-manno-oligosaccharide uptake and metabolism in *Bifidobacterium adolescentis* DSMZ 20083 from whole-genome microarray analysis": Dupplementary Figure 4

**Supplementary Figure 4**

**(A)**

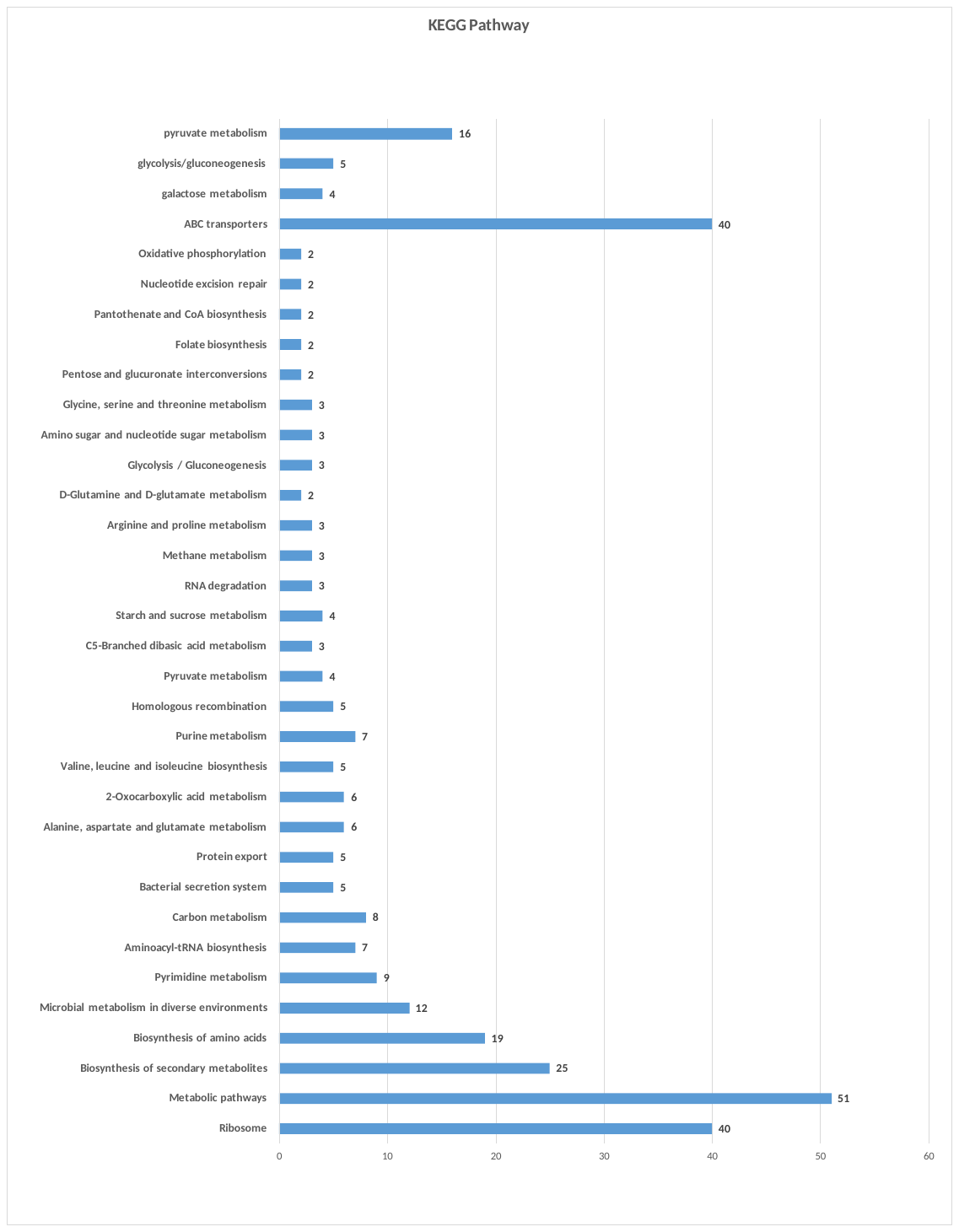

**(B)**

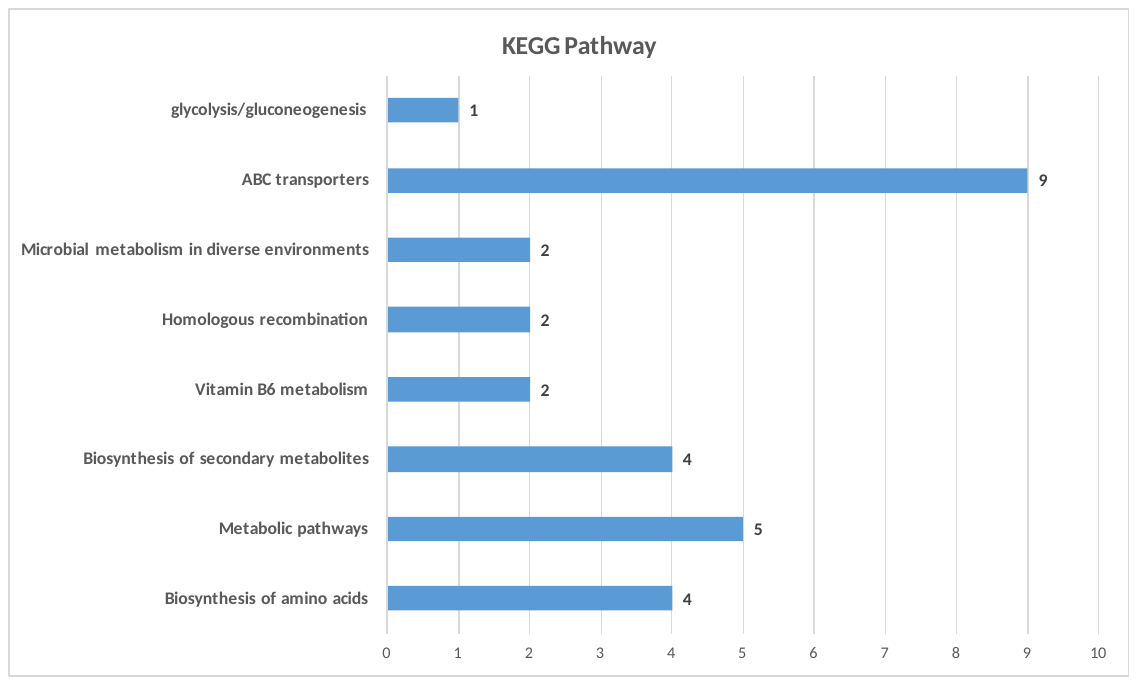
Figure Legend: KEGG enrichment analysis of (A) up-regulated genes (B) Down-regulated genes
