## Supplementary Figure 5 for "Insights into β-manno-oligosaccharide uptake and metabolism in *Bifidobacterium adolescentis* DSMZ 20083 from whole-genome microarray analysis"

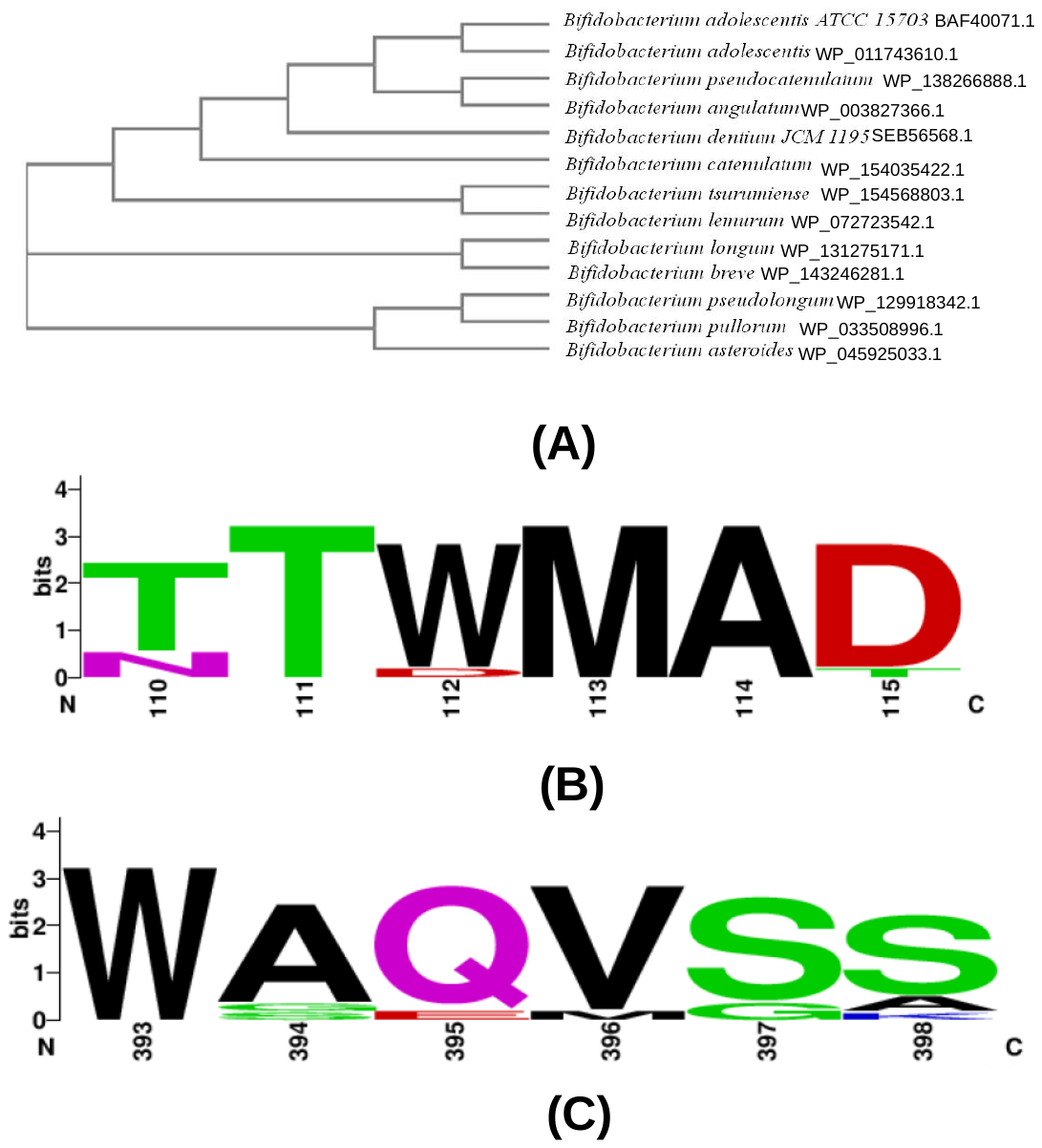
**Supplementary Figure 5**

0.01

**Supplementary Figure 5 Phylogenetic relationship and degree of conservation** of bglE (BAD_RS06835) with other ESBP homolouges from *Bifidobacteria* sp. **(A)** Phylogenetic tree based on multiple sequence alignment. **(B)** and **(C)** Sequence logo conservation of the bglE amino acid residues participating in H-bond formation with mannobiose.
