## Supplementary Table 1 for "Insights into β-manno-oligosaccharide uptake and metabolism in *Bifidobacterium adolescentis* DSMZ 20083 from whole-genome microarray analysis"

| **S.No.** | **Gene number** | **Forward primer** | **Reverse primer** |
| --- | --- | --- | --- |
| **1** | BAD_RS06080 | ACAAACGGGGGTATCCATCT | CGCTTTGTTGCGTACGAGT |
| **2** | BAD_RS06835 | TGAGGAAGCCATTTTTCAAAG | CCTTGCTTGAATCGATGGTC |
| **3** | BAD_RS06830 | CCTCGGGACAAGAAAGACAA | GGGGAATCAGCATGAATACG |
| **4** | BAD_RS06825 | AAGCAACAAGCAGTCGCTCT | ACAGGAACGGGAAGATCCAG |
| **5** | BAD_RS06820 | ATCCGCCATACTTGGAACG | ATGGCGTCGAAGGTGTAATC |
| **6** | BAD_RS06085 | CTCCCGTAAGTTCGGCTTC | GGAACCAACCCAGAAGTTCA |
| **7** | BAD_RS06090 | GCACCTGTCCTTCACGAAAT | TGTGGAACTTCGGATACCAG |
| **8** | gal E | GCCAGACCAAGCTGTTCG | CCGGATCCTCAAGTTCCAC |
| **9** | galM | CGGACAGCAATTTTCCATTT | GGGGAGACGATGATGTCCT |
| **10** | 16S rRNA | GAGCGAACAGGATTAGATAC | TCTTTGAGTTTTAGCCTTGC |

**Supplementary Table 1 Primer sequences of selected genes used in q-RT-PCR studies.**
