## Supplementary Table 2 for "Insights into β-manno-oligosaccharide uptake and metabolism in *Bifidobacterium adolescentis* DSMZ 20083 from whole-genome microarray analysis"

| **S.No.** | **Carbon source (0.1 % w/v)** | **Optical density (A_600nm_)** |
| --- | --- | --- |
| 1 | Negative control | 0.045 |
| 2 | Positive control | 0.572 |
| 3 | Guar gum | ND |
| 4 | Locust bean gum | ND |
| 5 | Konjac glucomannan | ND |
| 6 | Copra meal | ND |
| 7 | CM2 | 0.09 |
| 8 | CM3 | 0.34 |
| 9 | CM4 | 0.17 |
| 10 | DP2 GG-β-MOS | 0.16 |
| 11 | DP3 GG-β-MOS | 0.65 |
| 12 | DP4 GMOS | 0.13 |

**Supplementary Table 2** *In vitro* fermentation of β-mannans and pure β-manno-oligosaccharides by *B. adolescentis* DSMZ 20083

Negative control: SDM devoid of carbon source; Positive control: SDM containing 0.1% (w/v) glucose; ND: no growth. Pure GG-β-MOS (Mary *et al.* 2019) and GMOS (Srivastava *et al.* 2017) were prepared as per our previous studies. Briefly, guar gum and locust bean gum were hydrolyzed using GH26 endo-mannanase ManB-1601. Thereafter the precipitated and concentrated oligosaccharides were purified by size exclusion chromatography on Bio-gel P2. The fractionated oligosaccharides were freeze-dried and used in fermentation experiments.
